# Horizontal transfer of a LINE-RTE retrotransposon among parasite, host, prey and environment

**DOI:** 10.1101/2024.11.24.625053

**Authors:** T. Brann, F. S. de Oliveira, A. V. Protasio

**Author notes:** Emails: Toby Bran –, Fernanda Souza de Oliveira -, Anna V. Protasio –.

## Abstract

**Background:** Horizontal transfer of transposable elements is both impactful, owing to the subsequent transposition burst, and insightful, providing information on organisms’ evolutionary history. In eukaryotes, horizontal gene transfer (HGT) often involves transposable elements (TEs), host-parasite relationships, aquatic environments or any of them combined. The flatworm *Schistosoma mansoni* is a human parasite with two free-living aquatic stages (intercalated between a definitive human host and intermediate snail host) and has a sizable TE content. We aimed to identify and characterise potential instances of HGT leveraging new genomic resources available.

**Results:** Using the latest chromosome-scale genome assembly and available TE sequences we identify two putatively horizontally transferred elements, named Perere-3 and Sr3, in the *S. mansoni* genome. We demonstrate the presence of these TEs in the genomes of *Schistosoma* spp. intermediate hosts, most likely explained by HGT. Perere-3 / Sr3 were also found across a wide range of additional organisms not susceptible to schistosome infection, including turtles, fish and other molluscs.

**Conclusions:** We propose that the patchy distribution of Perere-3/Sr3 across the phylogenetic tree is best explained by HGT. This phenomenon is likely linked to the parasitic nature of schistosomes, as several snail species sharing the elements are susceptible to infection. However, presence of Perere-3/Sr3 in species beyond this relationship may suggest wider ancestral *Schistosomatidae* host ranges and/or undescribed schistosomes.

## Background

Horizontal gene transfer (HGT), unlike vertical genetic inheritance, refers to DNA acquisition via a non-parental genome. Although examples are found in all realms, HGT in eukaryotes occurs at a rate approximately 80 times less than in prokaryotes (Van Etten and Bhattacharya 2020). This is due in part to additional physical barriers present in eukaryotes, such as the nuclear envelope, compartmentalisation of the germline, where the transferred genes must be present to be propagated to future generations, and, in many metazoans, a protective skin or dermis (Kurland, et al. 2003).

Host-parasite relationships, which reduce or circumvent physical distances and barriers between organisms, are overrepresented in instances of eukaryotic HGT (Gilbert, et al. 2010). Parasites live for varied, but often extended, periods of time in direct contact or proximity to their hosts, often feeding on host tissue. This intimate host-parasite relationship provides an excellent environment for exposure to the other organism’s DNA. Another common medium for DNA “sharing” is aquatic environments. In them, many organisms such as algae, corals and fish, share and sample the same body of water, favouring HGT events (Abe, et al. 2020; Pérez-Etayo, et al. 2020).

Transposable Elements (TEs) are genetic entities capable of mobilisation within genomes. They are more frequently implicated in horizontal transfer events (referred to as “horizontal transfer of TEs” or HTT) than “traditional” host genes due to their inherent capacity to encode enzymes that mediate their excision and insertion into genomes. TEs are often present in many copies and can occupy large fractions of host genomes, such as 52% in Humans (Tang, et al. 2018), and 78% in Antarctic Krill (Shao, et al. 2023).

TE mobilisation has consequences in all genomes, disrupting or altering host genes and providing a substrate for exaptation or structural rearrangements (Young, et al. 2020). To mitigate the potential disruptive effects of TEs, organisms have evolved TE suppression mechanisms, including heterochromatin formation of TE-derived DNA preventing transcription and/or post-transcriptional silencing that degrades nascent TE mRNAs (Slotkin and Martienssen 2007). Foreign TEs acquired by HTT initially do not encounter any inherent resistance, resulting in a “transposition burst”: a rapid increase in TE copy number in the genome. Such a burst can change genome size, interrupting and shuffling genes and skewing GC content. For example, the horizontally transferred LINE-RTE BovB constitutes 18% of the cow (*Bos taurus*) genome and is also present in other mammals such as the platypus (*Ornithorhynchus anatinus*) and multiple lineages of snake (Walsh, et al. 2013; Puinongpo, et al. 2020). BovB dissemination was likely mediated by parasitic ticks with a broader range of ancestral hosts than currently observed (Ivancevic, et al. 2013). Examples such as this offer valuable insights into the evolutionary relationships and natural history of implicated species.

With approximately 40% of its genome derived from TEs, *Schistosoma mansoni*, a human parasitic flatworm and the best characterised species of the *Schistosomatidae* family, is considered TE-rich (International Helminth Genome Consortium 2019). These parasites have complex life cycles alternating between asexual reproduction in snails and sexual reproduction in vertebrate hosts (such as humans) with two short-lived aquatic stages: miracidia and cercariae (Nelwan 2019). Adult schistosomes can survive in their vertebrate hosts for many years or decades, the oldest recorded survival of individual worms being 37 years (Chabasse, et al. 1985). Parasite persistence in the snail host is variable but lifecycle completion requires at least 5 weeks in laboratory conditions (Kapp, et al. 2003).

The theoretical basis of *S. mansoni* involvement in HTT seems plausible given that the parasite has multiple aquatic life stages, extended periods of host association and TEs constitute at least one third of its genome (International Helminth Genome Consortium 2019), characteristics known to promote HTT (Gilbert, et al. 2010; Abe, et al. 2020; Pérez-Etayo, et al. 2020). In fact, an HTT event from salmonid to schistosome was proposed based on analysis of an EST library and subsequent amplification of salmonid-specific transposon-like sequences in *Schistosoma* DNA / cDNA (Melamed, et al. 2004). This work was later disproved on the basis that the level of salmon DNA contamination in the EST library by salmon was much higher than expected, potentially due to salmon sperm DNA being used as a DNA carrier during DNA isolation (Grunau and Boissier 2010; Wijayawardena, et al. 2015). Other claims of HTT in *Schistosoma* include transfers of MHC sequences from mice or rabbits (Imase, et al. 2001; Zhao, et al. 2009), DNA transposons from fish and frogs (Leaver 2001), receptor sequences from mammals (Hu, et al. 2003) and albumin from mice (Williams, et al. 2006; DeMarco, et al. 2007) have equally been contested based on DNA contamination, inadequate sampling of taxa, sequence conservation or incomplete / fragmented genome assemblies (Wijayawardena, et al. 2015). More recently, the transfer of a functional *cki* homolog in parasitic flatworms (including *S. mansoni*) from an unidentified metazoan has been proposed (Wendt and Collins III 2024), perhaps representing one of the few (as yet) undisputed cases of HGT.

Recently, a chromosomal-level genome assembly for *S. mansoni* (Buddenborg, et al. 2021) and high quality genomes of Schistosoma snail hosts (Nong, et al. 2022; Young, et al. 2022; Bu, et al. 2023) have been made available, opening the opportunity for a comprehensive investigation into horizontal transfer in a medically relevant organism with a high potential for gene transfer. Here, we leverage these genomic resources to identify and characterise instances of HTT between *S. mansoni* and its hosts and wider metazoans implicated in this phenomenon.

## Methods

For a preliminary assessment of horizontal transfer in selected genomes (Table 1), we used a library of available and curated *S. mansoni* TEs (Supplementary Table ST1, Supplementary File SFile1) as a query in a BLASTN search (v2.12.0) (Camacho, et al. 2009) with predefined “relaxed” parameters that improve the identification of contiguous TE sequences in species where they may have diverged and/or be highly fragmented in relation to the query sequences used (Galbraith, et al. 2020). These parameters are ‘`-reward 3 -penalty -4 -xdrop_ungap 80 -xdrop_gap 130 -xdrop_gap_final 150 -word_size 14 -dust yes -gapopen 30 -gapextend 6 -outfmt 6 -evalue 1e-5’` and define what we, from now on call, a “relaxed BLASTN”. In addition to published and available TEs (Drew and Brindley 1997; Drew, et al. 1999; Laha, et al. 2002; Arkhipova, et al. 2003; Copeland, et al. 2003; DeMarco, et al. 2004; Feschotte 2004; Laha, et al. 2004; Copeland, et al. 2005; DeMarco, et al. 2005; Laha, et al. 2005; DeMarco, et al. 2006; Jacinto, et al. 2011), we manually curated Perere-3 and Sr3 consensuses (Supplementary File SFile2) using the previously published sequences (DeMarco, et al. 2005; Laha, et al. 2005; Venancio, et al. 2010) and published methodology (Goubert, et al. 2022) with the *S. mansoni* genome (Buddenborg et al. 2021, PRJEA36577, WormBase ParaSite release WBPS19).

**Table 1 -.** Genome accessions used in preliminary identification of horizontally transferred elements. Organisms were selected for their relationship, either evolutionarily or parasitically to *Schistosoma mansoni*, as described in “Notes”.

| Organism | Assembly | Notes |
| --- | --- | --- |
| *Schistosoma mansoni* | GCA_000237925.5 | Parasite |
| *Trichobilharzia regenti* | GCA_944472135.2 | Other *Schistosomatidae* |
| *Heterobilharzia americana* | GCA_944470555.2 |  |
| *Fasciola hepatica* | GCA_948099385.1 |  |
| *Clonorchis sinensis* | GCA_003604175.2 | Other trematodes |
| *Bulinus truncatus* | GCA_021962125.1 | Intermediate host |
| *Biomphalaria straminea* | GCA_021533235.1 |  |
| *Lymnaea stagnalis* | GCA_900036025.1 |  |
| *Homo sapiens* | GCF_000001405.40 | Definitive |

We used the compiled TE library with manually curated Perere-3 and Sr3 to annotate the *S. mansoni* genome (Howe, et al. 2017; Buddenborg, et al. 2021) available in WormBase ParaSite release WBPS19 (PRJEA36577), using RepeatMasker v4.1.236 (Smit, et al. 2013) with default parameters except for `‘-no_is -gff -s -a’`. To investigate conserved protein domains in consensus sequences of Perere-3 and Sr3, we retrieved open reading frames with EMBOSS’ `‘getorf -minsize 500’` (Rice, et al. 2000) from their DNA sequences and submitted them to InterProScan5 (Jones, et al. 2014). Percentage identities of consensus and relevant domains (L1-EN - cd09076 and RT-Pol - PS50878) were generated using Jalview’s Pairwise Alignment (Waterhouse, et al. 2009). When evaluating Perere-3/Sr3 genomic insertions, these were considered “Complete” when conserved protein domains “Exo_endo_phos” and “RVT_1” were both present in at least 99% of their full length, as identified with Pfam (Mistry, et al. 2021).

To study the relationship between Perere-3 and Sr3 insertions within the *S. mansoni* genome, we retrieved copies with a minimum length of 3,200bp and used them to construct a distance tree with iqtree (v2.2.3) (Minh, et al. 2020) with default parameters except for `‘--seqtype DNA -T AUTO --alrt 1000’.` The resulting tree was visualised in ITOL (v5) (Letunic and Bork 2021) and Perere-3 and Sr3 coloured to highlight segregation. We calculated relative TE quantities directly from our output of RepeatMasker (see above) and represented these using the `‘treemapify’` (v2.5.5) and `‘ggplot2’` (v3.4.3) (Wickham 2011) packages, implemented in R Studio (Build 554) (RStudio Team 2020). To investigate substitution rates of these elements with respect to their consensus sequences, we collected Kimura values (with divCpGMod) from RepeatMasker’s alignment file generated using the `‘-a’` option and represented them in a histogram.

To further assess TE activity, we used the *terminal branch length* (TBL) method, previously used for analysis of zebrafish TEs (Chang, et al. 2022). Fragments greater than 500bp were aligned with `‘mafft —-auto’` (Katoh, et al. 2005), trimmed with TrimAl (Capella-Gutiérrez, et al. 2009) with the `‘-gt’` parameter set to 0.01. Trees were then constructed per TE using `‘FastTree -nt -gamma’` (Price, et al. 2010) and TBLs of the element fragment tree were then extracted using `‘termlength.py’`.

To provide a framework for our proposed TE transfer among *Schistosoma* and selected species, we constructed a species tree using sequences obtained with reciprologs (https://github.com/glarue/reciprologs, with default parameters), an approach that identifies “reciprocal best hits” *i.e.* amino acid sequences that are pairwise best hits across all species sampled. These reciprologs genes were then aligned per-gene using `‘mafft —-auto’` (Katoh, et al. 2005) and subsequently concatenated to generate a single alignment per species of all reciprocal best hits. Species level phylogeny was then constructed with iqtree (Minh, et al. 2020) with parameters `‘--seqtype AA -T AUTO --alrt 1000 -B 1000’`. For the input into reciprologs, we obtained trematode amino acid sequences from WormBase ParaSite (release WBPS19) and filtered them to include just the longest isoform of every gene. For the snail genomes, for which amino acid sequences were not readily available, BUSCO (Simão, et al. 2015) was used (with default parameters and the `‘mollusca_obd10’` database) to identify putative conserved amino acid sequences derived from the mollusc genomes. The resulting amino acid sequences were then used as inputs for reciprologs. Further to our identification of Perere-3/Sr3 sequences in non-*S. mansoni* genomes, we proceeded to manually curate (Goubert, et al. 2022) their consensus sequences in these genomes (Table 2 and Table 3). We introduced a variation on the first step of the manual curation (see protocol by Goubert et al. 2022), replacing the stringent BLASTN search with our “relaxed BLASTN” approach to account for greater sequence differences between query (*S. mansoni* Perere-3 and Sr3) and subject.

**Table 2 -.** Genome accessions used in construction of *Schistosoma* phylogeny. All genomes used are available at WormBase Parasite release 19. *Trichobilharzia regenti* was used as an outgroup for analysis.

| Organism | Assembly | Accession |
| --- | --- | --- |
| *Plakobranhus ocellatus* | PoB_v1 | GCA_019648995.1 |
| *Schistosoma mansoni* | v10 | GCA_000237925.5 |
| *Schistosoma japonicum* | ASM636876v1 | GCA_025215515.1 |
| *Schistosoma turkestanicum* | tdSchTur1.1 | GCA_944470395.2 |
| *Schistosoma rodhaini* | tdSchRodh2.1 | GCA_944470435.2 |
| *Schistosoma spindale* | tdSchSpin1.1 | GCA_946903255.1 |
| *Schistosoma margrebowiei* | tdSchMarg1.1 | GCA_944470205.2 |
| *Schistosoma mattheei* | tdSchMatt1.1 | GCA_944470405.2 |
| *Schistosoma intercalatum* | tdSchInte2.1 | GCA_944470385.2 |
| *Schistosoma haematobium* | tdSchHaem2.1 | GCA_944470465.2 |
| *Schistosoma guineensis* | tdSchGuin1.1 | GCA_944470375.2 |
| *Schistosoma curassoni* | tdSchCurr1.1 | GCA_944474815.3 |
| *Schistosoma bovis* | tdSchBovi2.1 | GCA_944470445.2 |
| *Trichobilharzia regenti* | tdTriRege1.1 | GCA_944472135.2 |

**Table 3 -.** Genome accessions used in construction of molluscan phylogeny. All genomes used are available on NCBI Genome List. *Plakobranchus ocellatus* was used as an outgroup for analysis.

| Organism | Assembly | Accession |
| --- | --- | --- |
| *Plakobranthus ocellatus* | PoB_v1 | GCA_019648995.1 |
| *Achatina immaculata* | ASM976088v1 | GCA_009760885.1 |
| *Arion vulgaris* | ASM2079622v1 | GCA_020796225.1 |
| *Cepaea nemoralis* | Cnem_1.0 | GCA_014155875.1 |
| *Candidula unifasciata* | ASM976088v1 | GCA_905116865.2 |
| *Physella (Physa) acuta* | ASM2847654v2 | GCA_028476545.2 |
| *Lymnaea stagnalis* | v1.0 | GCA_900036025.1 |
| *Radix auricularia* | ASM207201v1 | GCA_002072015.1 |
| *Ampullaceana balthica* | Rbalt genome v1 | GCA_944989445.1 |
| *Bulinus truncatus* | Btru.v1 | GCA_021962125.1 |
| *Anisus vortex* | AniVort1.1 | GCA_949126835.1 |
| *Biomphalaria straminea* | Bstr_hk_v1 | GCA_021533235.1 |
| *Biomphalaria glabrata* | BioGlab47.1 | GCF_947242115.1 |
| *Biomphalaria pfeifferi* | UNM_Bpfe_1.0 | GCA_030265305.1 |

To investigate the relationship between consensus sequences of Perere-3 and Sr3-like elements from Schistosoma and snail genomes, we calculated their pairwise nucleotide distances (including that of Sr2 as outgroup) using EMBOSS’ `‘distmat’` and the Jukes-Cantor substitution model (Rice, et al. 2000). These relationships are represented using the `‘pheatmap’` package (Kolde and Kolde 2015) to generate a hierarchical clustering tree.

To assess any potential transfer of Perere-3 and Sr3 across metazoa, we constructed a dendrogram of combined novel Perere-3 / Sr3-like sequences from *Schistosoma* and molluscan organisms (Table 2 and 3) and LINE_RTE sequences from RepBase (Bao, et al. 2015). Sequences were aligned with `‘mafft –auto’` and trimmed with TrimAl (Capella-Gutiérrez, et al. 2009) with `‘-gt’` set to 0.25. This dendrogram was generated with iqtree using default parameters except for `‘-bb 1000 -alrt 1000’`. To identify the extent of transfer of Perere-3 and Sr3 across metazoa, we scanned 3446 high-quality genome assemblies (Scaffold N50 > 1Mbp, accessed from GenBank using the esearch function, August 2023) (Benson, et al. 2018) using our “relaxed BLASTN” approach. The species list was filtered for entries with available speciation data (n=2271) at timetree.org (Kumar, et al. 2022). We manually annotated the tree to indicate “taxa by phyla” for all non-chordates and “taxa by class” for chordates. Some clades representing a single or small numbers of species were not annotated to improve clarity of figure.

To explore the species implicated in the metazoan transfer, we highlighted species with either the highest “Max Length Hit”, “Percentage Identity” or “Number of hits over 1,000bp”. We also included hits from notable taxa - the only bird, mammal and a coral (highest of the seven corals identified) and *Schistosoma japonicum* (the only *Schistosoma* organism with speciation data). To explore horizontal transfer across these metazoan species we first manually curated Perere-3 / Sr3-like TEs for all organisms in Table 5 and generated both a species phylogeny and TE consensus tree for comparison. Species phylogeny for organisms in Table 5 was generated with timetree.org (Kumar, et al. 2022) and TE consensus tree constructed with iqtree and `‘-bb 1000 -wbt -alrt 1000’`parameters after alignment with `‘mafft —-auto’`.

**Table 4 -.** Full length copies of Perere-3 and Sr3 are found across *Schistosoma spp*. Perere-3 and Sr3 copies identified using the ‘relaxed BLASTN’ and manually curated *S. mansoni* Perere-3 and Sr3 consensus sequences. All genomes used are available on WormBase ParaSite, with accessions listed in Table 2.

| Species | Perere-3 |  |  | Sr3 |  |  |
| --- | --- | --- | --- | --- | --- | --- |
|  | % Genome | n(>2,900bp) | Max Full Length | % Genome | n(>2,900bp) | Max Full Length |
| *S. japonicum* | 3.81 | 977 | 3239 | 3.88 | 982 | 3242 |
| *S. turkestanicum* | 5.98 | 1156 | 3225 | 5.36 | 1135 | 3241 |
| *S. rodhaini* | 5.31 | 1563 | 3241 | 5.81 | 1574 | 3241 |
| *S. mansoni* | 4.91 | 1073 | 3241 | 5.43 | 1079 | 3244 |
| *S. spindale* | 5.5 | 1482 | 3237 | 6.04 | 1486 | 3252 |
| *S. margrebowiei* | 4.8 | 891 | 3257 | 5.35 | 903 | 3257 |
| *S. mattheei* | 5.1 | 1277 | 3247 | 5.65 | 1284 | 3287 |
| *S. intercalatum* | 5.13 | 1253 | 3256 | 5.69 | 1257 | 3256 |
| *S. haematobium* | 4.81 | 1511 | 3253 | 5.32 | 1514 | 3269 |
| *S. guineensis* | 4.94 | 1130 | 3253 | 5.49 | 1134 | 3268 |
| *S. curassoni* | 4.96 | 1168 | 3259 | 5.51 | 1177 | 3266 |
| *S. bovis* | 4.93 | 1115 | 3249 | 5.46 | 1122 | 3249 |

**Table 5.**
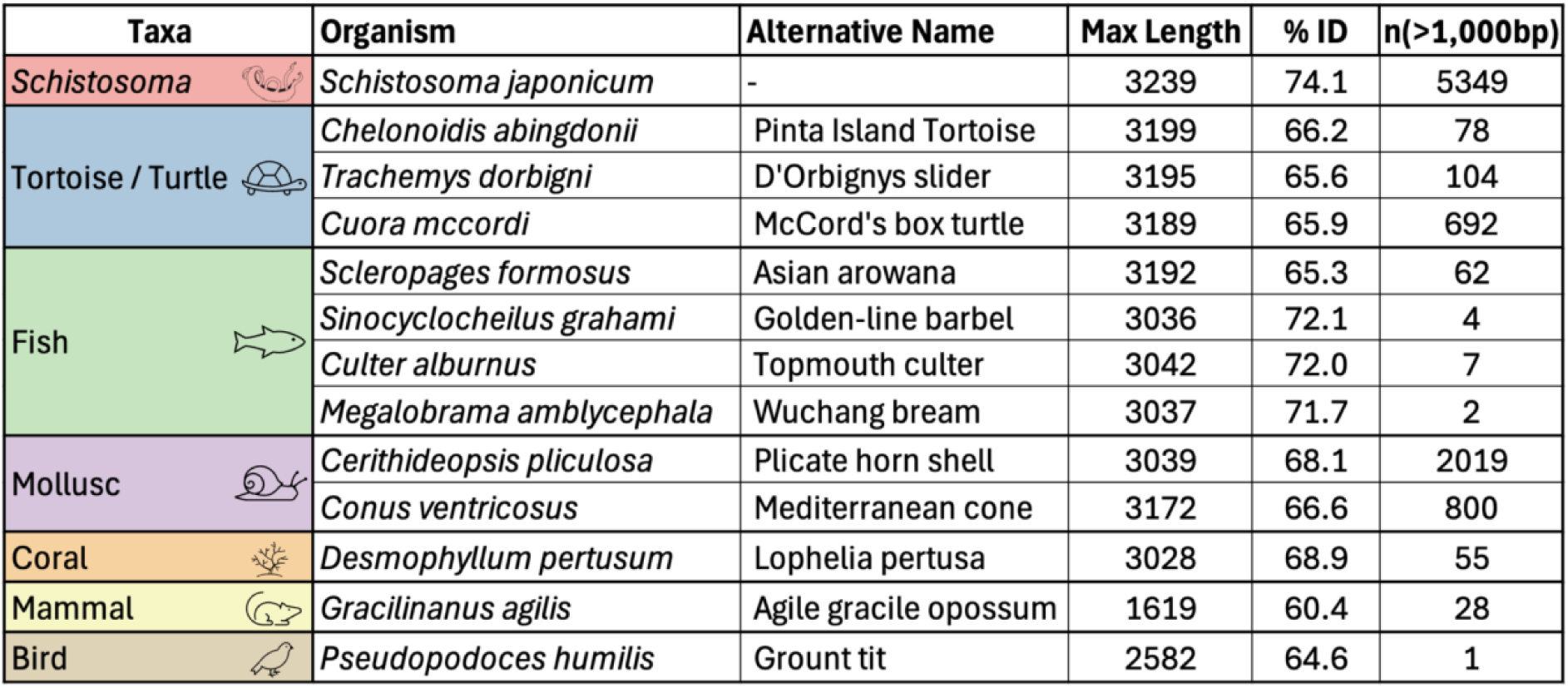
Select metazoan species identified from Figure 4. Species were chosen based on the highest maximum length, highest percentage identities and highest n(>1,000bp) found. In addition to these, notable taxa identified were also included (coral, mammal and bird).

| Taxa | Organism | Alternative Name | Max Length | % ID | n(>1,000bp) |
| --- | --- | --- | --- | --- | --- |
| *Schistosoma* | *Schistosoma japonicum* | - | 3239 | 74.1 | 5349 |
| Tortoise / Turtle | *Chelonoidis abingdonii* | Pinta Island Tortoise | 3199 | 66.2 | 78 |
|  | *Trachemys dorbigni* | D'Orbignys slider | 3195 | 65.6 | 104 |
|  | *Cuora mccordi* | McCord's box turtle | 3189 | 65.9 | 692 |
| Fish | *Scleropages formosus* | Asian arowana | 3192 | 65.3 | 62 |
|  | *Sinocyclocheilus grahami* | Golden-line barbel | 3036 | 72.1 | 4 |
|  | *Culter alburnus* | Topmouth culter | 3042 | 72.0 | 7 |
|  | *Megalobrama amblycephala* | Wuchang bream | 3037 | 71.7 | 2 |
| Mollusc | *Cerithideopsis pliculosa* | Plicate horn shell | 3039 | 68.1 | 2019 |
|  | *Conus ventricosus* | Mediterranean cone | 3172 | 66.6 | 800 |
| Coral | *Desmophyllum pertusum* | Lophelia pertusa | 3028 | 68.9 | 55 |
| Mammal | *Gracilinanus agilis* | Agile gracile opossum | 1619 | 60.4 | 28 |
| Bird | *Pseudopodoces humilis* | Grount tit | 2582 | 64.6 | 1 |

All processing, analysis and visualisation scripts used in this publication are available on GitHub at: https://github.com/tbrann99/Schisto_HTT/.

## Results

### Finding *Schistosoma* TE-like sequences in the parasite’s intermediate hosts

We explored the possibility of HTT in *Schistosoma mansoni* using a relaxed BLASTN search of known *S. mansoni* TEs against other Schistosomatidae (*Trichobilharzia regenti*, *Heterobilharzia americana*) and trematodes (*Fasciola hepatica, Clonorchis sinensis*) as well as intermediate (*Bulinus truncatus, Biomphalaria straminea, Lymnaea stagnalis*) or definitive hosts (*Homo sapiens*). Using this approach, we found significant similarity to two LINE-RTEs known as Perere-3 and Sr3 in the three genomes corresponding to Schistosoma intermediate hosts, *i.e.* molluscs (Figure 1). Prospective TEs are present at near-full-length of the *S. mansoni* consensus sequence used as query in the search (91.3% and 90.8% in *Bu. truncatus,* 98.3% and 97.9% in *Bi. straminea*, 91.3% and 90.8% in *L. stagnalis* for Perere-3 and Sr3 respectively), suggesting similarity is not simply due to the presence of highly conserved TE domains but instead represents a full-length match. Additionally, in these organisms, many near-full-length (more than 90% full length, approximately 2,900bp for Perere-3 and Sr3) copies were identified, indicating transposition of these elements. In contrast, few “Other TEs” were identified above the e-value filter in intermediate snail hosts. We did not detect significant hits of Perere-3/Sr3 in the other trematodes closely related to *Schistosoma*, namely *C. sinensis* or *F. hepatica*. As these trematodes are evolutionarily closer to *S. mansoni* than the molluscan intermediate host, the presence of Perere-3 and Sr3 is unlikely to be a product of vertical inheritance. This first observation led us to hypothesise that a HTT event between molluscs and their parasites could have occurred, either directly or indirectly.

**Figure 1 -.**
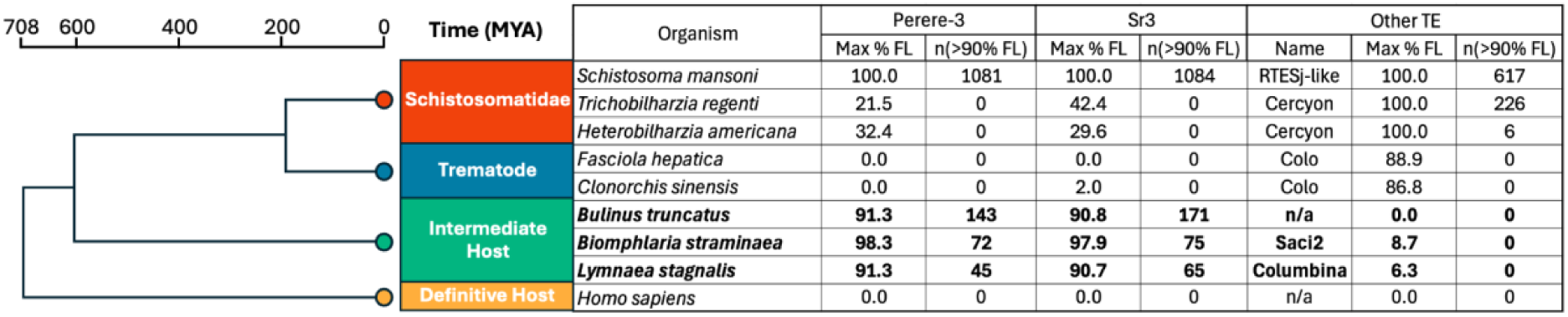
The genome of the intermediate host of *Schistosoma mansoni* contains *S. mansoni*-like TEs. Select organisms with varying relationships to *S. mansoni* were scanned with a relaxed BLASTN to identify the presence of *S. mansoni* TEs in these genomes. “Max % FL” represents the highest percentage of the *S. mansoni* TE consensus sequence that was mapped to the genome. “n(90% FL) was calculated as the number of copies identified over 90% of the full length of the respective elements. The highest non-Perere-3 or non-Sr3 element (“Other TE”) is also shown to highlight potential differences between Perere-3 / Sr3-derived results and other elements. Simplified phylogeny (left) was extracted from timetree.org (Kumar, et al. 2022), representative genomes used for each group highlighted with *.

### Perere-3 and Sr3 in the *Schistosoma mansoni* genome

We characterised Perere-3 and Sr3 in *S. mansoni*. In the first instance, we took the sequences of Perere-3 and Sr3 previously described by Venancio et al. (2010) and used the most recent version of the *S. mansoni* assembly to update their consensus sequences using a manual curation approach that better represent the range of TE insertions found in the genome. The resulting consensus sequences for Perere-3 and Sr3 are 3,207 and 3,209 base pairs respectively (Supplementary File SFile2). These elements have high sequence similarity: at the DNA level, they share a nucleotide percentage identity of 79.81%. Both elements encode full-length L1-endonuclease (L1-EN) and the reverse transcriptase (RT Pol), for which their amino acid percentage identities are 83.81% and 82.59% related to one another respectively (Figure 2A).

**Figure 2 –.**
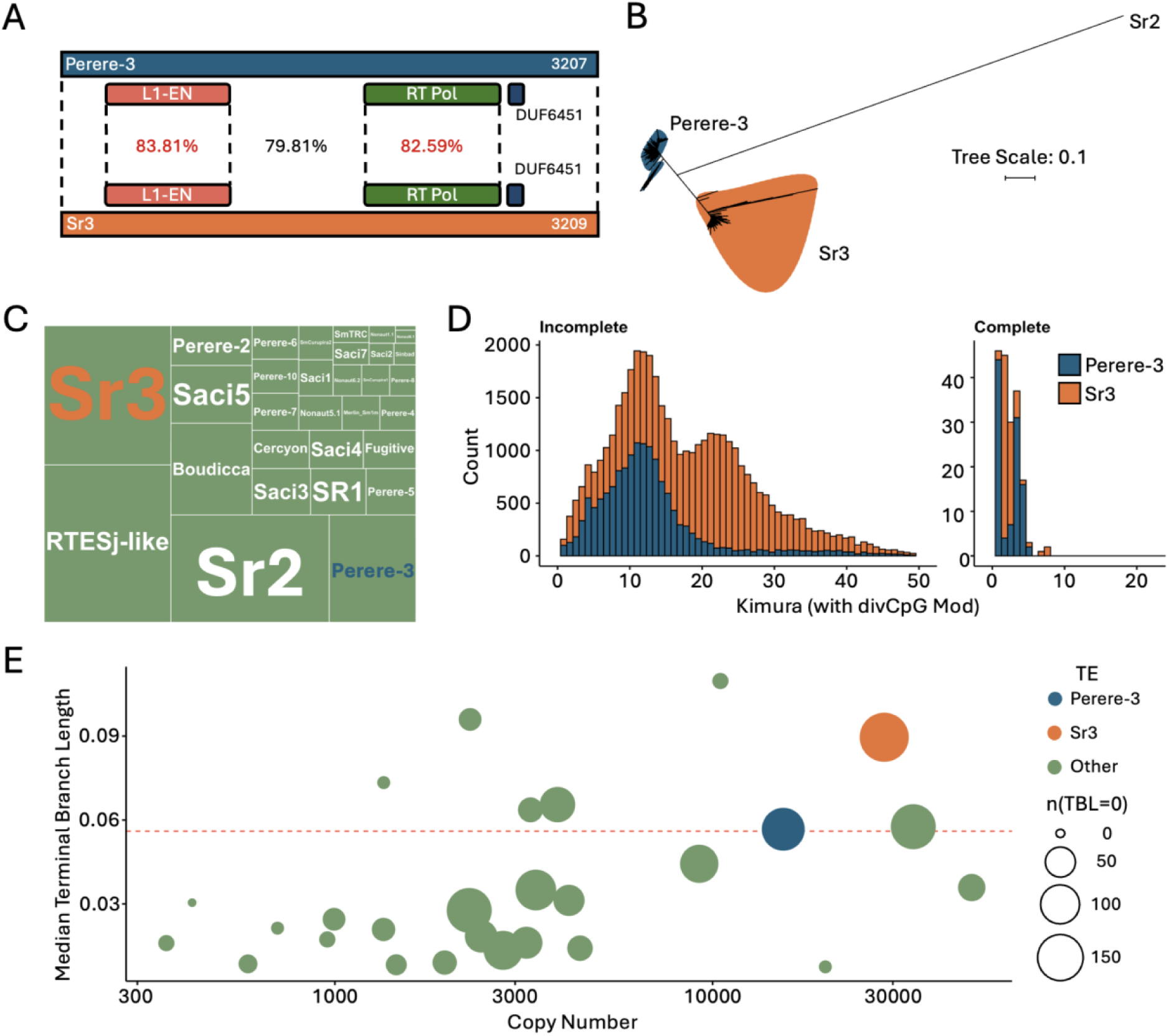
Perere-3 and Sr3 are similar LINE-RTEs with different transposition histories within the *S. mansoni* genome. **A)** Protein domain topology of Perere-3 and Sr3, with percentage identity (DNA) of the entire element (in black) and the L1-EN and RT Pol protein domains (in red). **B)** Tree of full-length (>2,900bp) TE insertions in the *S. mansoni* genome with Sr2 LINE-RTE as outgroup. **C)** Genomic coverage of TEs of the *S. mansoni* genome. Total coverage with TE library of 24.7%. **D)** Kimura profile of Perere-3 and Sr3 in the *S. mansoni* genome, divided into “Incomplete” and “Complete” elements, where complete indicates at least 99% of both the L1-EN and RT Pol domains. **E)** *Terminal branch length* (TBL) analysis of all *S. mansoni* TE copies, assessing copy number against median terminal branch length. The size of the datapoint indicates the number of instances where the TBL was 0. Red dashed line represents the mean terminal branch length of all elements.

Despite their close sequence similarity, these TEs are considered to transpose independently from one another. To confirm this, we retrieved from the genome all Perere-3 and Sr3 copies with a minimum length of 3,200bp and built a sequence similarity dendrogram (Figure 2B). Our results show that sequences arising from Perere-3 or Sr3 TE transposition events segregate into two distinct clusters. Furthermore, the relative distance between these two LINE-RTEs and others from the same organism suggests that Perere-3 and Sr3 may have relatively recently shared a common TE ancestor (Figure 2B). Differential transposition activity has resulted in these elements having different proportions of genomic coverage: Perere-3 and Sr3 occupy 2.1% and 4.0% of the genome respectively (Figure 2C). Further evidence for their independent activity can be seen in their profile of accumulated mutations represented in the differential kimura distributions (Figure 2D). Kimura distances represent the evolutionary distance of a given insertion from the consensus sequence from which relative age may be inferred. Our data show that Sr3 has a distinct bimodal distribution of kimura values, with a peak at both 12 and 22, in comparison to Perere-3 which is unimodal with a single peak corresponding to a kimura value of 12. This indicates differential bursts of transposition between these elements and pointing to independent transposition.

To further understand the dynamics of Perere-3 and Sr3 in *S. mansoni*, particularly given their potential involvement in horizontal transfer events, we aimed to assess ongoing transposition activity of these elements. Based on their completeness, these elements are capable of transposing: of the 43,800 Perere-3/Sr3 annotated fragments, 432 (approximately 1%) have at least 99% of the length of the protein domains required for TE transposition, namely the endonuclease and reverse transcriptase domains. However, final proof of ongoing or very recent transposition would be given by presence of identical TE insertions. To this end, we use a TBL approach which provides an estimation of the relative age of all genomic TE copies (Chang, et al. 2022), with low TBLs indicating more recent transpositions. We demonstrate that Sr3 and Perere-3 have 173 and 124 copies with a TBL of 0 (Figure 2E), indicating many identical pair-wise matches, and providing evidence of recent enough transposition events that did not have time to accumulate mutations.

### Evolutionary Spread and Activity of Perere-3 and Sr3 Across Schistosoma and Molluscan Hosts

In the first section, we shown that Perere-3 and Sr3 are found extensively across the *Schistosoma* genus but not in full length nor in high numbers in *Trichobilharzia*, a member of the *Schistosomatidae* family, or wider Trematoda (Figure 1). We broadened our search beyond the *S. mansoni* genome to evaluate the presence and extent of transposition across the *Schistosoma* genus and to more accurately determine the evolutionary timing of a potential transfer event. We searched for Perere-3/Sr3 sequences in 11 additional *Schistosoma* genomes and found that all contained full copies of the TEs (Table 4).

Our analysis of TBL across the *Schistosoma* genus (Figure 3A) suggests that these elements have been active throughout speciation. Whilst similar numbers of near full length (>2,900bp) elements are observed across *Schistosoma*, Perere-3/Sr3 from *S. japonicum* and *S. turkestanicum* have higher TBLs medians and fewer near 0 TBLs, indicating that a higher proportion of transposition events in these organisms are older than in those species most recently diverged (Figure 3A). We found a significant increase in terminal branch length between *S. japonicum* and *S. turkestanicum* and the newer diverged schistosomes (Kruskal-Wallis and Dunn’s test, *p<0.001*). The widespread presence and activity of these elements across *Schistosoma* indicate that the horizontal transfer event likely occurred in a common ancestor of the genus.

**Figure 3 -.**
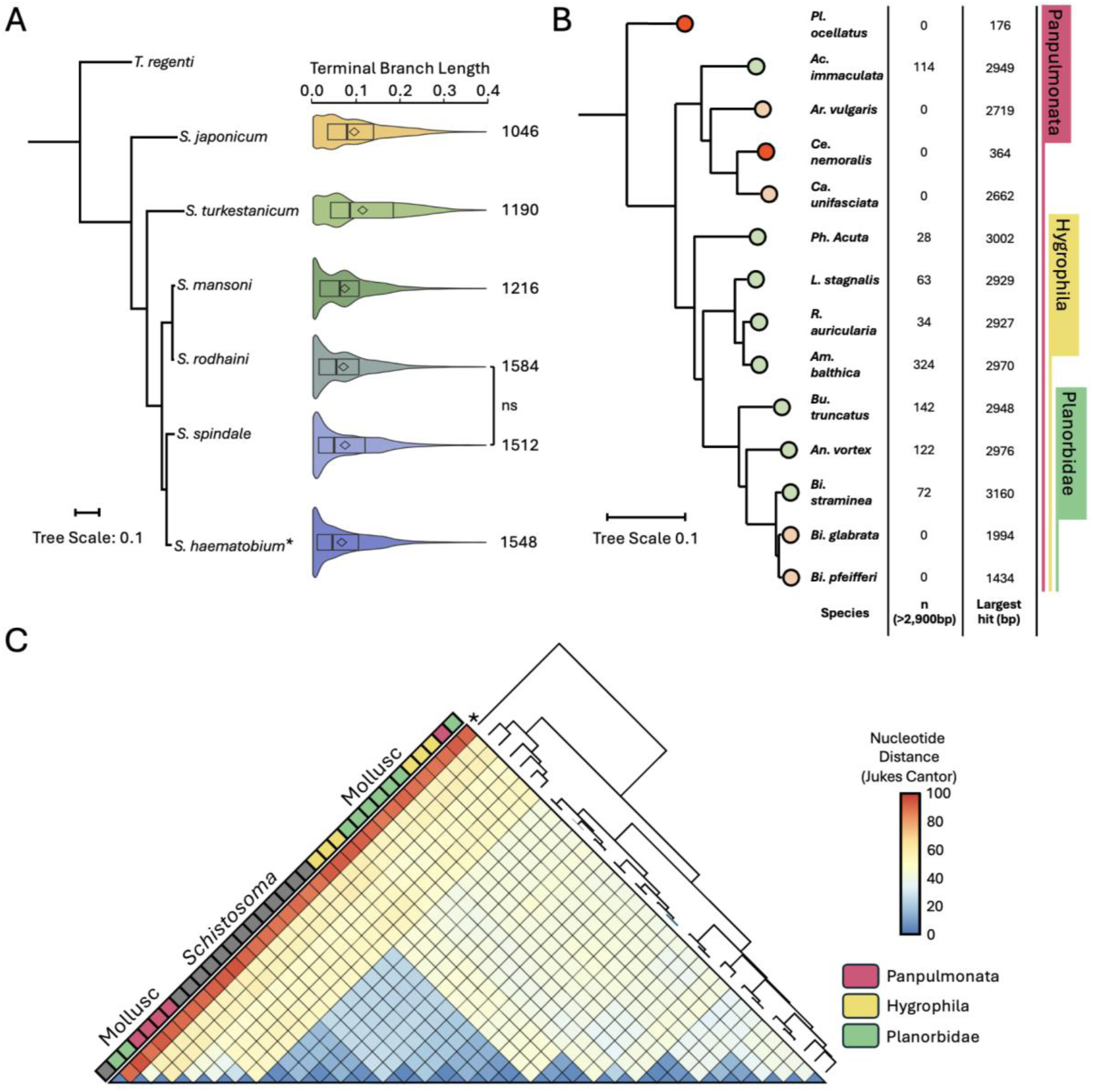
*S. mansoni* Perere-3 and Sr3-like sequences are found across *Schistosoma sp.* and molluscs, including snails susceptible to infection. **A)** *Schistosoma* species tree and terminal branch length (TBL) distribution of Perere-3 and Sr3-like insertions aligned to said tree. Species tree was constructed from reciprocal orthologs across all species. *Schistosoma* phylogeny was constructed with all species available in WormBase ParaSite. *S. mattheei*, *S. intercalatum*, *S. haematobium*, *S. guineensis*, *S. curassoni* and *S. bovis* were collapsed into a single clade representing the *haematobium* group (*), in line with previous phylogenetic assessment of *Schistosoma (Leger and Webster 2017)*. *Trichobilharzia regenti* was used to root the tree, though no full-length Perere-3 / Sr3-like sequences were identified in this organism (Figure 1). TBL values plotted correspond to the *S. haematobium* genome. Diamonds indicate the mean. Testing for terminal branch length differences between species with a Kruskal-Wallis test and Dunn’s post-hoc analysis demonstrated significance (*p<0.001*) across all pairwise-species comparisons except *S. spindale* and *S. rodhaini* (*p>0.05*). Number of elements greater than 2,900bp is shown to the right of the violin plot. **B)** Rooted species tree of Panpulmonata, including Planorbidae snails, the intermediate hosts of *Schistosoma spp.* and related Hygrophila snails. *Plakobranchus ocellatus* was used as an outgroup. Molluscan genes were obtained with BUSCO (see Methods). Leaves are colour-coded to indicate presence of elements over 2,900bp (green), over 1,000bp (orange) or shorter (red). **C)** Heatmap representing distance matrix (DNA) of manually curated TE consensus sequences from *Schistosoma* and panpulmonate snails alongside hierarchical clustering tree. Snails are labelled Planorbidae (green), Hygrophila (yellow) or panpulmonate (pink). *S. mansoni* LINE-RTE Sr2 was added as an outgroup, as indicated by the ‘*’ in the tree.

Having found extended presence of Perere-3 and Sr3 in a wider range of *Schistosoma spp*, we wanted to test whether these TEs are also present in the respective intermediate hosts (class: Planorbidae) and other molluscs across Panpulmonata. We use the ‘relaxed BLASTN’ approach with *S. mansoni* Perere-3 and Sr3 as queries on the genomes of selected Planorbidae and Panpulmonata and build a corresponding species tree to represent species relatedness (Figure 3B). We found significant BLASTN hits corresponding to more than 90% of the full length of *S. mansoni* Perere-3 and Sr3 consensus sequences (green in Figure 3B, Supplementary Table ST2) in 8 molluscan species. For instance, *Am. balthica* contains 324 copies over 2,900bp in length. Two of the three species of *Biomphalaria* (namely *B. glabrata* and *B. pfeifferi*) have a marked reduction in number of copies as well as their maximum length, suggesting TEs in these species are no longer active and the remaining identifiable copies have accumulated many mutations.

To explore the relationship between *Schistosoma* and molluscan Perere-3 and Sr3-like sequences, we manually curated these TEs from the 12 molluscan genomes for which significant BLASTN hits were found. To find out how closely related these sequences are to each other, we built a nucleotide distance matrix (Figure 3C) of the *Schistosoma* and molluscan TEs consensus sequences. A hierarchical clustering approach was used to identify intersequence relationships and we found that *Schistosoma* Perere-3 and Sr3-like sequences cluster in the centre of the heatmap within the curated molluscan TEs (Grey, Figure 3C). The high similarity observed between *Schistosoma* and molluscan Perere-3 and Sr3-like elements and positioning of the *Schistosoma* clade within a continuum of molluscan TE sequences, further adds to the evidence of a horizontal transfer event. Additionally, we found Perere-3 / Sr3 in Hygrophila snails, which are not Planorbidae and are not known to be susceptible to *Schistosoma* infection.

### Assessment of Perere-3 and Sr3 across metazoa

To assess the relationship between *Schistosoma*/molluscan Perere-3 and Sr3-like sequences and the wider LINE-RTE superfamily of TEs, we extracted consensus sequences of LINE-RTEs from the RepBase database (Bao, et al. 2015) and used these to construct a sequence similarity dendrogram. Our analysis included two well-known horizontal TE transfers of LINE-RTEs namely BovB (Ivancevic, et al. 2013) and AviRTE (Suh, et al. 2016). We show that Perere-3 and Sr3 sequences from *Schistosoma* and gastropoda cluster tightly together and, at the same time, separately from the known TE transfers, indicating a distinct transfer event from those previously described (Figure 4A). Several LINE-RTEs from RepBase cluster around our curated Sr3 and Perere-3 sequences, such as those from *Crassostrea gigas* (Pacific Oyster), *Aplysia californica* (Californian Sea Hare, a sea slug), *Saccoglossus kowalevskii* (Acorn Worm, a free-living, hemichordate), *Danio rerio* (Zebrafish) and *Chrysemys picta bellii* (Western Painted Turtle).

**Figure 4 -.**
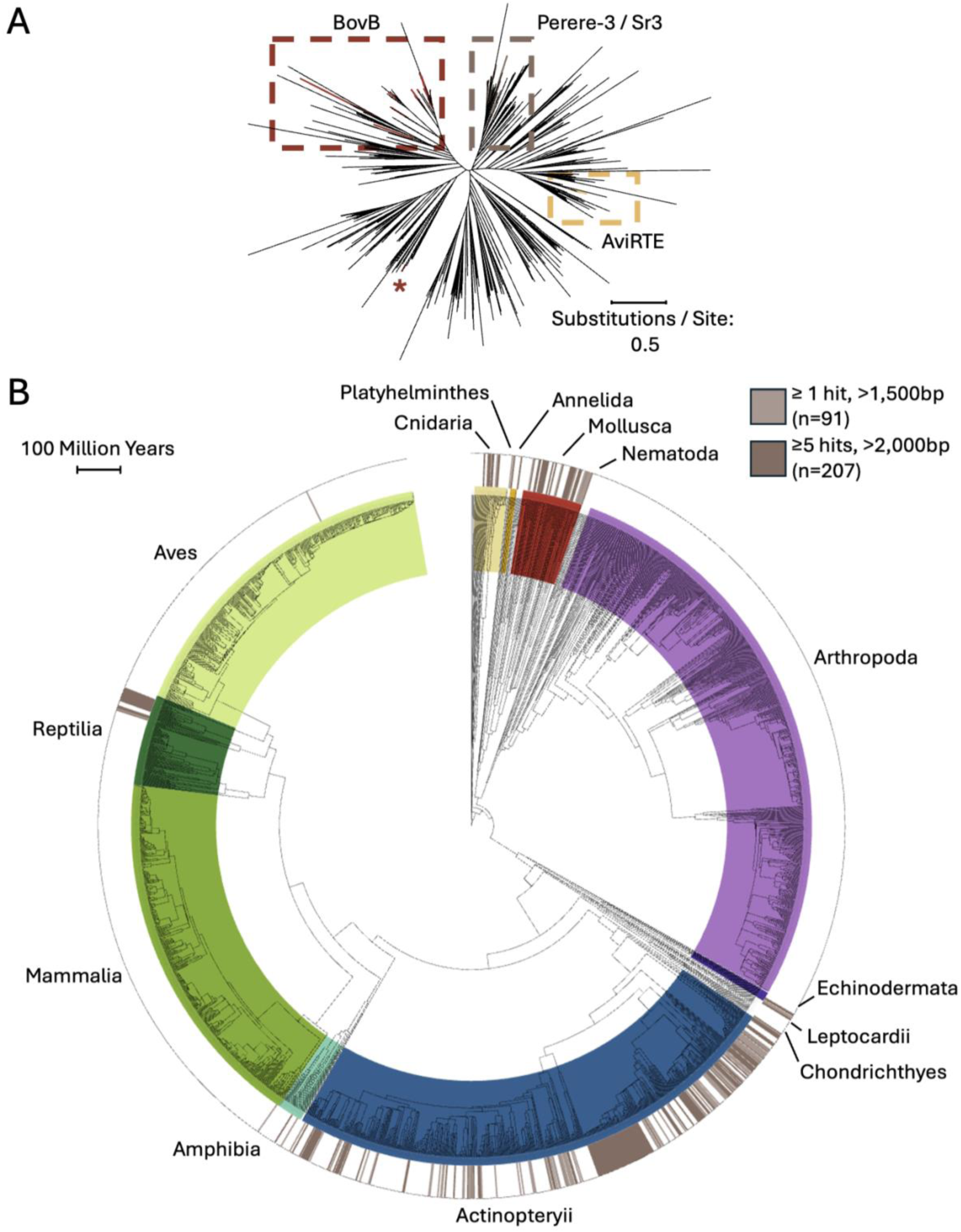
Perere-3 and Sr3 are found across metazoa. **A)** Dendrogram of consensus sequences of LINE-RTEs found in RepBase plus *Schistosoma* and mollusc Perere-3 / Sr3-like sequences, highlighting known horizontally transferred groups BovB (Ivancevic, et al. 2013) and AviRTE (Suh, et al. 2016). * Indicates additional BovB sequence that did not cluster within other BovB sequences. **B)** Presence of Perere-3 / Sr3-like sequences across metazoan genomes available at Timetree.org (Kumar, et al. 2022). Relevant hits were classified as either weak (light brown: at least one hit > 1,500bp) or strong (dark brown: at least one hit > 2,000bp and at least five hits > 1,000bp). Perere-3 and Sr3 were used as a query, results shown for the TE generating higher results, TEs were plotted separately in Supplementary Figure SF1.

We therefore hypothesised that Perere-3 and Sr3 may exist outside *Schistosoma* and molluscs and, as RepBase is limited to user-submitted, previously curated TEs, we extended our search to all available, high quality metazoan genomes. This was done with the relaxed BLASTN approach using *S. mansoni* Perere-3 and Sr3 as queries against all metazoan genomes (contig N50 > 1Mbp) available in GenBank (n=3,446, Supplementary Table ST3). An organism was determined to be a “weak” candidate for HTT if it returned a blast hit >1,500bp, while “strong” candidate required at least one larger than 2,000bp and at least five hits >1,000bp. Stringency around the number of hits identified helps to differentiate between contamination in the genome assembly resulting in a single hit and *bona fide* hits, which would have many copies. Using these criteria we found, out of the 3,446 genomes tested, 117 organisms to be “weak” candidates for Perere-3/Sr3 HTT while 256 of these were “strong” candidates. To visualise the presence/absence of Perere-3/Sr3-like sequences in these species and their respective relationships, we used timetree.org and filtered for genomes for which speciation data was available in this database (n=2,271). From this subset of genomes, 207 and 91 were “weak” and ‘strong’ hits respectively.

We found Perere-3/Sr3-like sequences discontinuously across the metazoan tree (Figure 4B), including reptiles, such as the previously mentioned *Chrysemys picta bellii*, amphibians, marine invertebrates, molluscs, cnidaria, a single bird and a single mammal and more extensively across fish. Such a diverse number of taxa from which these hits derived suggests that Perere-3 and Sr3 have been extensively transferred across metazoans. Given the similarity of Perere-3 and Sr3 (Figure 2A) and the expected nucleotide distances of such TEs, highly similar results were seen for Perere-3 and Sr3 when searched against metazoan genomes (Supplementary Figure SF1).

Of the 298 organisms identified as containing Perere-3 / Sr3-like sequences, we highlight several species (Table 5) which were selected either due to having large numbers of hits over 1,000bp, highest single largest hits or being notable taxonomic clades. To investigate TEs and species relatedness, we curated Perere-3 and Sr3-like sequences in these species. We first evaluated their sequence similarity and presence of conserved protein domains and found that, despite a large evolutionary distance across taxa (Figure 5A), all but two species of curated TEs encode full length L1-end and RT domains and shared the overall TE domain topology (Figure 5B, Supplementary Figure SF2). We then compared a dendrogram of curated consensus sequences to the corresponding species tree. We show extensive phylogenetic incongruence between the species and TE trees (Figure 5). For example, despite fish and *S. mansoni* diverging more than 600 million years ago (Figure 5A), curated Perere-3 and Sr3-like sequences from fish are closely related to their *S. mansoni* counterparts, contradicting what would be expected by vertical inheritance. Such phylogenetic incongruence between species for sequences sharing such levels of similarity suggests extensive horizontal transfer of Perere-3 and Sr3, beyond *Schistosoma* and mollusca and across metazoa.

**Figure 5.**
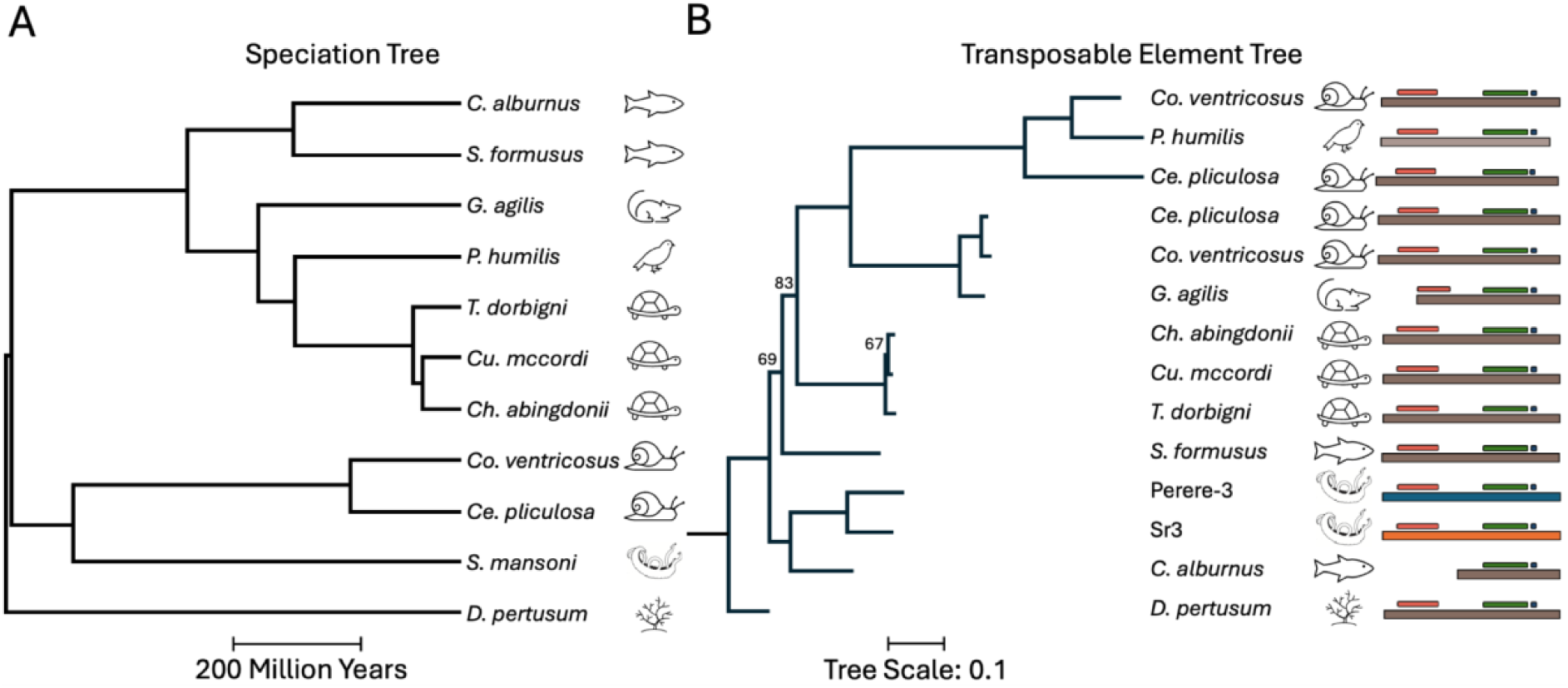
Juxtaposition of TE and species trees highlights the phylogenetic incongruence between them and is evidence in support of the horizontal transfer of Sr3 and Perere-3. **A)** Speciation tree generated with timetree.org showing divergence times of selected organisms. **B)** Tree of manually curated TE consensus sequences of Perere-3 and Sr3 from the same organisms in A. Two distinct elements were curated for *Conus ventricosus* and *Cerithideopsis pliculosa*. *Culter alburnus* and *Gracilinanus agilis* curated elements were 5’ truncated. Ultrafast bootstrap values lower than 98 are shown. TE domains of curated elements are represented with boxes to the right of the panel; L1-EN (red), RT_Pol (green) and DUF-6451. Protein domains of all consensus sequences were mapped using the methodology for Figure 2A.

## Discussion

Our preliminary investigation using BLASTN on the genomes of trematodes and *Schistosoma* hosts (namely human and Planorbidae snails) with *S. mansoni* TEs, revealed that two elements, Perere-3 and Sr3, are well represented in the genomes of *Schistosoma spp.* intermediate hosts (*Lymnaea stagnalis*, *Bu. truncatus*, *Bi. straminea*), while no other element shares the same degree of similarity among this group. Perere-3 and Sr3 were absent from the definitive human host and from the non-*Schistosoma* trematode genomes sampled, suggesting either loss in the trematode lineages sampled or a later gain in *Schistosoma*. Owing to the parasitic relationship and prolonged contact between *Schistosoma* and their intermediate host, we hypothesised that Perere-3 and Sr3 could represent examples of horizontally transferred TEs. Previously documented horizontal transfer events involving *S. mansoni, e.g.* with Salmonids (Melamed, et al. 2004), Humans (Yu, et al. 2008) and Sturgeon (Hale, et al. 2010) have all subsequently been refuted (Wijayawardena, et al. 2015). As these rare transfer events can provide insights into the evolutionary history of organisms and their relatedness (Ivancevic, et al. 2013; Daubin and Szöllősi 2016) we aimed to further characterise and confirm our initial observation of HTT of Perere-3 / Sr3 and examine evidence for the direction of transfer.

Our first focus was on assessing Perere-3 and Sr3 within the *S mansoni* genome. Perere-3 and Sr3 are two LINE-RTEs with high sequence similarity that exist at high frequency in the *S. mansoni* genome. With a percentage identity of 79.81% across their full-length sequence, these elements fall just outside the “80:80:80 rule” (Wicker, et al. 2007) which justifies their classification as separate TE families (Figure 2A). We demonstrate that the diversity of near-full length insertions of Perere-3 and Sr3 in the *S. mansoni* genome segregated into two distinct clusters (Figure 2B). We propose that these elements were at one point a single element that underwent extensive transposition and divergence.

Within the *S. mansoni* genome, Perere-3 and Sr3 represent the second and fourth most abundant TEs, respectively (Figure 2C). Together, these elements comprise 6% of the *S. mansoni* genome and account for approximately one-fifth of the organism’s TE content, indicating extensive past transposition. In addition to their considerable abundance, we show that these elements are capable of transposition, as indicated by 432 insertions having full-length (Figure 2D) conserved L1-EN and RT protein domains, the two enzymes needed for active transposition. In fact, these TEs are likely actively transposing, as evidenced by the 430 copies with terminal branch lengths of 0 (Figure 2E), *i.e.* not even one nucleotide difference between the pairs of insertions considered. While by no means a confirmation of horizontal transfer, historical transposition bursts and periods of ongoing activity suggest lack or reduced capability of sequence-specific resistance by the host genome, making these TEs good candidates of HTT (Gilbert and Feschotte 2018). Perere-3 and Sr3’s transposition activity and high abundance in *S. mansoni* has contributed to its increased genome size, providing a large source of homologous substrate for recombination, which may have contributed in turn to the expansion of gene families in the TE-rich subtelomere of *S. mansoni* (Brann, et al. 2024).

We further investigated the presence or absence of Perere-3 and Sr3 across *Schistosoma* and Panpulmonata lineages by manually curating consensus sequences of these elements in selected species. We were able to identify and curate Perere-3 and Sr3-like elements from representative *Schistsosoma* lineages having constructed a phylogeny for all genome assemblies available at the time of writing. We show that the most recently diverged Schistosoma species (*S. haematobium, S. spindale, S. mansoni* and *S. rodhaini*) have a larger proportion of active elements (reflected in the lower TBL median) than the more basal species *S. japonicum* and *S. turkestanicum* (Figure 3A). We cannot directly attribute speciation of *Schistosoma* to the horizontal acquisition of an element in the last common ancestor, but such an event is likely to have had a large impact on the genus’ evolution, interrupting genes and / or genic regions or promoting structural rearrangements. Venancio et al. (2010) hypothesised the transposition of Perere-3 and Sr3 contributed to the speciation of African schistosomes. To confirm this, ‘speciation genes’ associated with *Schistosoma* diversification would first need to be identified (Nosil and Schluter 2011) and the subsequent contribution of Perere-3 and Sr3 to these be investigated.

Our analysis identified significant hits of Perere-3 and Sr3-like sequences in Planorbidae genomes, a family of snails susceptible to *Schistosoma* infection. The large evolutionary distance that separates these two animal groups, approximately 600 million years (Kumar, et al. 2022), suggests a horizontal transfer event has occurred. To identify the direction of transfer, we extended our relaxed BLASTN to include other, related snails from Hygrophila and the wider Panpulmonata order and demonstrated the extended presence of these elements (Figure 3B). For instance, *Bu. truncatus*, susceptible to *S. japonicum* infection, and *Bi. straminea*, susceptible to *Schistosoma spp.* infection, have 142 (~ 1Mbp or 0.25% of the genome) and 72 copies above 90% of the full length of the consensus sequences of *S. mansoni* Perere-3 and Sr3. Given the multiple occurrences and completeness of copies found across molluscan genomes, such an observation is unlikely to result from DNA contamination during genome assembly, as has been proposed for previous putative *S. mansoni* HGT observations (Wijayawardena, et al. 2015). Interestingly, *Bi. glabrata* and *B. pfeifferi* had only degraded copies, despite their close relationship to *Bi. straminea* (Figure 3B), suggesting that these elements have stopped transposition sometime post-divergence from *Bi. straminea*.

To find further evidence in support of HTT and discern potential evolutionary relationships between TE–derived sequences in *Schistosoma* and panpulmonate, we calculated the pairwise nucleotide distance of manually curated, consensus sequences of Perere-3 and Sr3-like from species in these groups (Figure 3C). We show that *Schistosoma* Perere-3 and Sr3-like sequences cluster within the comparatively more diverse molluscan sequences. This observation suggests a potential route for the horizontal transfer of Perere-3 and Sr3 from panpulmonate to *Schistosoma* presumably via the parasitic relationship of an ancestral *Schistosoma* panpulmonate host-parasite pair. The two groups of molluscan sequences formed by the “insertion” of the *Schistosoma* cluster (Figure 3C) only loosely correspond to either the constructed phylogenetic tree or panpulmonates’ susceptibility to schistosome infection. This discordance could be explained by yet undescribed parasitic relationships between snail and host, or relationships that cannot occur due to a lack of geographic overlap rather than biological incompatibility. From the sequence and species relationships investigated here, we propose that Perere-3 and Sr3 LINE-RTEs, which have extensively transposed in *Schistosoma spp.*, could have been acquired via horizontal transfer from an ancestral snail host.

To investigate if these TEs could have been further transferred in more distantly related species, we expanded our search through a collection of metazoan genomes. In this approach we queried the genomes of 2,271 organisms (for which speciation data was available at timetree.org) for sequences with high similarity to *S. mansoni* Perere-3 or Sr3. We identified Perere-3/Sr3-like sequences across 298 organisms, 204 of which were classified as “strong” hits (more than five hits larger than 1,000bp and one more than 2,000bp), including reptiles, gastropods and fish (Figure 4B). A weak hit was identified from one mammal (*Gracilanus agilis*) and bird (*Pseudopodoces humilis*). These weak hits are likely contamination of the genome assembly, as the multiple large copies were not identified, indicating transposition had not occurred in these genomes. Other species had strong hits, indicative of not only presence of a single TE but TE activity. For example, the painted turtle *Chrysemys picta bellii*, which organisms from the superfamily *Schistosomatoidae* do infect turtles, and trematode infection has been described (Johnson, et al. 1998). Fish host a remarkable number of digenean parasites (Bullard and Overstreet 2008) and the horizontal transfer of genetic material between predator and prey has been documented via parasite intermediaries (Kambayashi, et al. 2022). Fish, such as *Oreochromis* identified here have been trialled in the biological control of schistosomiasis via predation of snail populations (Slootweg, et al. 1994) and will also eat free-living aquatic stages of the trematode life cycle (Orlofske, et al. 2015). It is possible that the feeding of fish, on either Perere-3 / Sr3-containing snails and schistosomes may provide an avenue by which HGT has occurred. Many of the identified snails are hosts for multiple parasites with complex life cycles, for example, *Lymnaea stagnalis* is “a cosmopolitan vector of trematodes infecting diverse vertebrates” (Gilbert, et al. 2010). It is possible that these panpulmonates act as a distribution centre for horizontally transferred material across different host-parasite combinations and life cycles that facilitate the extensive transfer of Perere-3 and Sr3 across metazoan organisms in and around aquatic environments. Water-filtering organisms such as sponges and corals are known for their ability to accurately reflect the metagenome of a given environment (Cai, et al. 2022). In aquatic environments where Perere-3 and Sr3 are actively expanding within and transferring between organisms, it seems likely that this phenomenon would extend to organisms sampling such an environment (Figure 6.)

**Figure 6.**
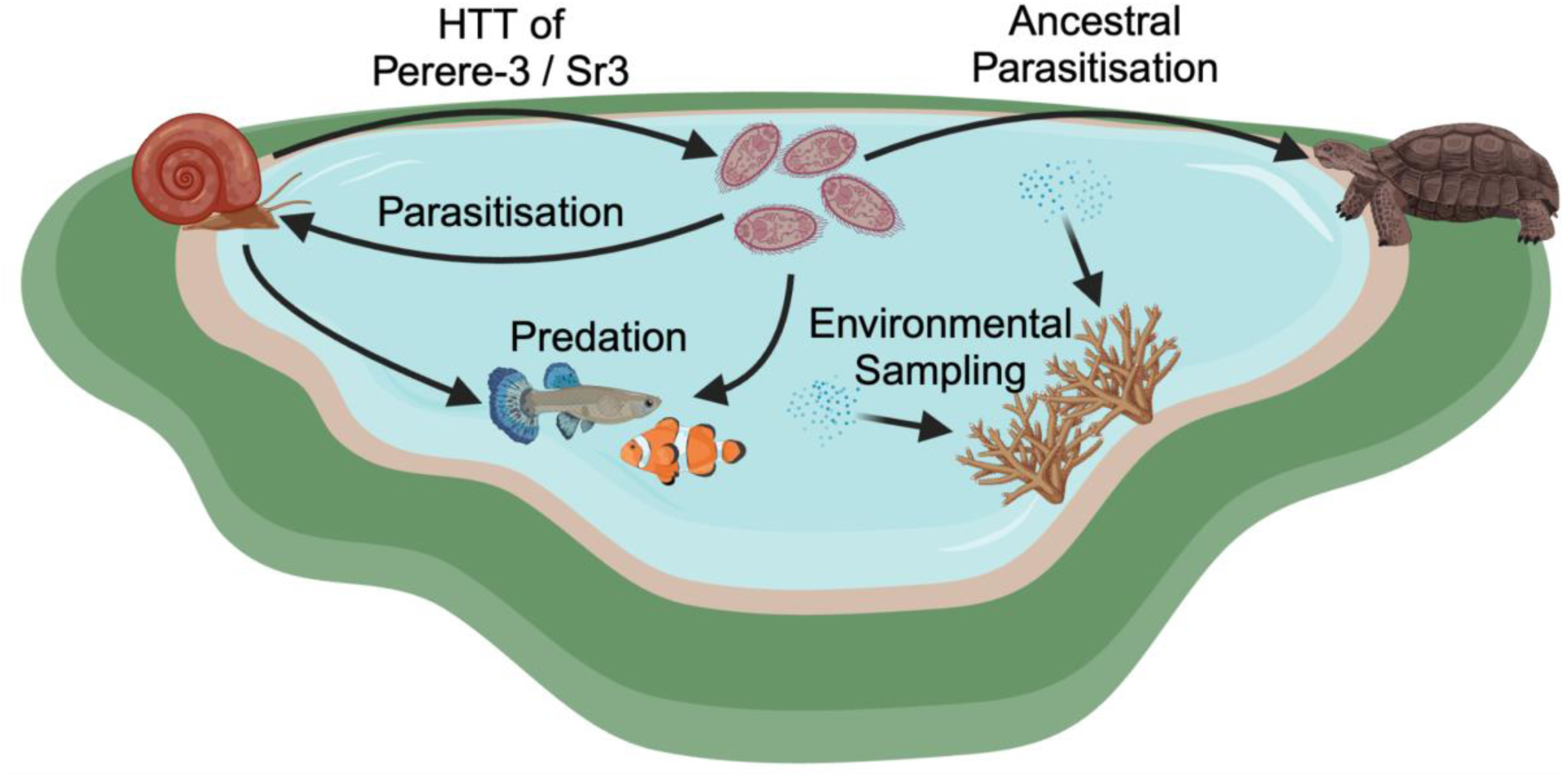
Proposed network of organisms implicated in the transfer of Perere-3 and Sr3. Organisms not to scale, arrows not representative of direction of Perere-3 and Sr3 transfer but of network interactions. Image generated using biorender.com.

The phylogenetic distribution of Perere-3/Sr3-containing species is patchy and cannot be explained by vertical inheritance alone (unless many simultaneous TE losses occurred in many genomes). This observation suggests that the horizontal transfer of Perere-3 and Sr3 extends well beyond the host-parasite relationships between Schistosoma and molluscs. A comparison of the speciation tree and the TE phylogeny generated for these organisms (Figure 5) demonstrates this phylogenetic incongruence.

## Conclusion

We were able to leverage available genomic resources for *S. mansoni* and the high-quality genomes of other organisms to identify Perere-3 and Sr3-like sequences and characterise the extent of horizontal transfer of these LINE-RTEs in a way not previously possible. We show evidence in support of horizontal transfer of Perere-3 / Sr3 from panpulmonate snails to *Schistosoma* parasites, which, in light of further evidence showing the spread of these sequences further through the phylogenetic tree, appears as the tip of the iceberg of a much broader transfer dynamic. Owing to the rebuking of previous *S. mansoni* transfer events, we believe this represents one of the few robust horizontal transfer examples available in *S. mansoni*, due to the number of organisms implicated, quality of genomes sampled, and breadth of organisms involved.

## Declarations

### Ethics approval and consent to participate

Not applicable.

### Consent for publication

Not applicable.

### Availability of data and materials

All processing, analysis and visualisation scripts used in this publication are available on GitHub at: https://github.com/tbrann99/Schisto_HTT/.

### Author Contributions

Study was conceived by TB. Methods and analysis were implemented by TB and AVP. Manual curation of Perere-3 and Sr3 was completed by AVP and manual curation of elements in *Schistosoma* (excluding *S. mansoni*) and snails was completed by FdSO. The manuscript was drafted by TB and AVP. All authors edited and approved the final version of the manuscript.

## Supporting information

SFile1

SFile2

## Acknowledgments

Dr James Galbraith, University of Exeter, for discussions around early observations. Dr Alex Suh, University of Bonn, for help and ideas around early drafts of analysis. Kristian Sorensen, for contributing to literature analysis, manuscript review and discussions on horizontal transfer context.Caroline Walker, University of Cambridge, for illustrative feedback of proposed network of organisms.

## Supplementary Information

<u>Supplementary Table ST1</u>. **Source of *Schistosoma mansoni* transposable element consensus sequences used in analysis.** Reported refers to the relevant citation in which the TE was identified / described and accessions of respective elements provided where consensus sequences were submitted.

<u>Supplementary Table ST2</u>. **Relaxed BLASTN output of Perere-3 and Sr3 on molluscan genomes used in analysis.** Results reported for Perere-3 and Sr3 separately. ‘% Genome’ refers to the percentage of the genome occupied by the respective element, ‘n(>2,900bp)’ the number of hits above 2,900bp and ‘Max Full Length’ the length of the largest hit identified.

<u>Supplementary Table ST3.</u> **Assembly information and relaxed BLASTN output for all genomes selected for analysis.** All metazoan genome assemblies with an N50 greater than 1,000,000bp were identified and searched for the presence of Perere-3 and Sr3 using the ‘Relaxed BLASTN’. Blast results for all species are shown for Perere-3 and Sr3 respectively, highlighting the percentage identity (%id), length and E-value for the best hit, as identified by bit score. Additionally, the number of hits over 1,000bp for the given element was also highlighted (n>1,000bp). For visualisation, a species phylogeny was constructed using timetree; species with an appropriate entry are highlighted in “Present in TimeTree” column.

<u>Supplementary Figure SF1</u>. **Separate Perere-3 (A) and Sr3 (B) blast results across high-quality metazoan genomes.** Presence of Perere-3 / Sr3-like sequences across metazoan genomes available at Timetree.org. Relevant hits were classified as either weak (light brown: at least one hit > 1,500bp) or strong (dark brown: at least one hit > 2,000bp and at least five hits > 1,000bp).

<u>Supplementary Figure SF2</u>. **Reverse transcriptase and transposable element consensus distance trees provide similar topology.** Trees of manually curated TE consensus sequences of Perere-3 and Sr3 from organisms of Table 4. Two distinct elements were curated for *Conus ventricosus* and *Cerithideopsis pliculosa*. *Culter alburnus* and *Gracilinanus agilis* curated elements were 5’ truncated. Reverse transcriptase domain tree generated using InterProScan5 to identify relevant protein domains. Both trees generated using `‘iqtree -bb 1000 -wbt -alrt 1000’` after alignment with mafft.

<u>Supplementary File SFile1</u>. ***Schistosoma mansoni*’s transposable element library used for analysis.** Compiled from sources listed in Supplementary Table 1.

<u>Supplementary File SFile2</u>. **Manually curated Perere-3 and Sr3 sequences from *Schistosoma mansoni*.** Sequences curated using methodology outlined by Goubert et al. 2022.

