## Supplementary material for "Horizontal transfer of a LINE-RTE retrotransposon among parasite, host, prey and environment": SF2

### Supplementary Figure 2

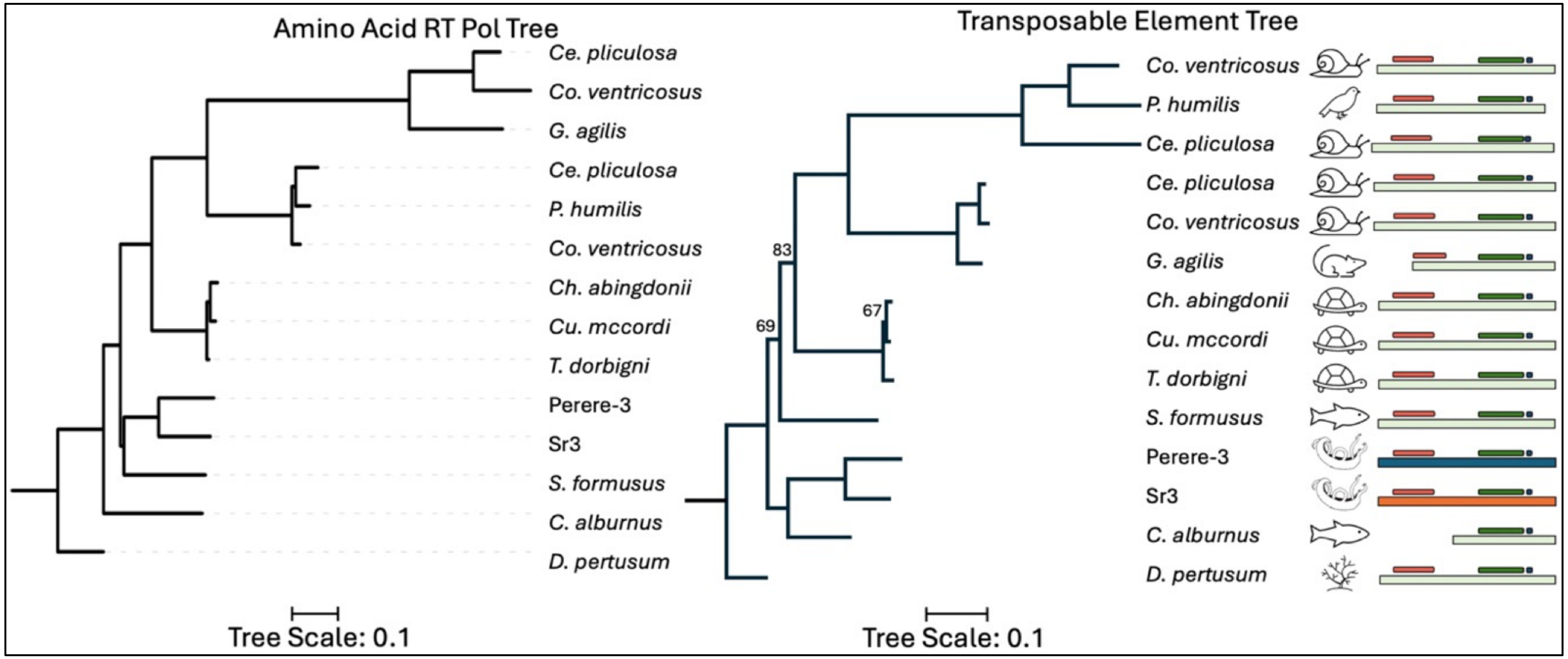


**Supplementary Figure 2 – Reverse transcriptase and transposable element consensus distance trees provide similar topology.** Trees of manually curated TE consensus sequences of Perere-3 and Sr3 from organisms of Table 4. Two distinct elements were curated for *Conus ventricosus* and *Cerithideopsis pliculosa. Culter alburnus* and *G. agilis* curated elements were 5' truncated. Reverse transcriptase domain tree generated using InterProScan5 to identify relevant protein domains. Trees generated using `'iqtree -bb 1000 -wbt -alrt 1000'` after alignment with `mafft`.
