## Supplementary material for "Horizontal transfer of a LINE-RTE retrotransposon among parasite, host, prey and environment": SF1

### Supplementary Figure 1

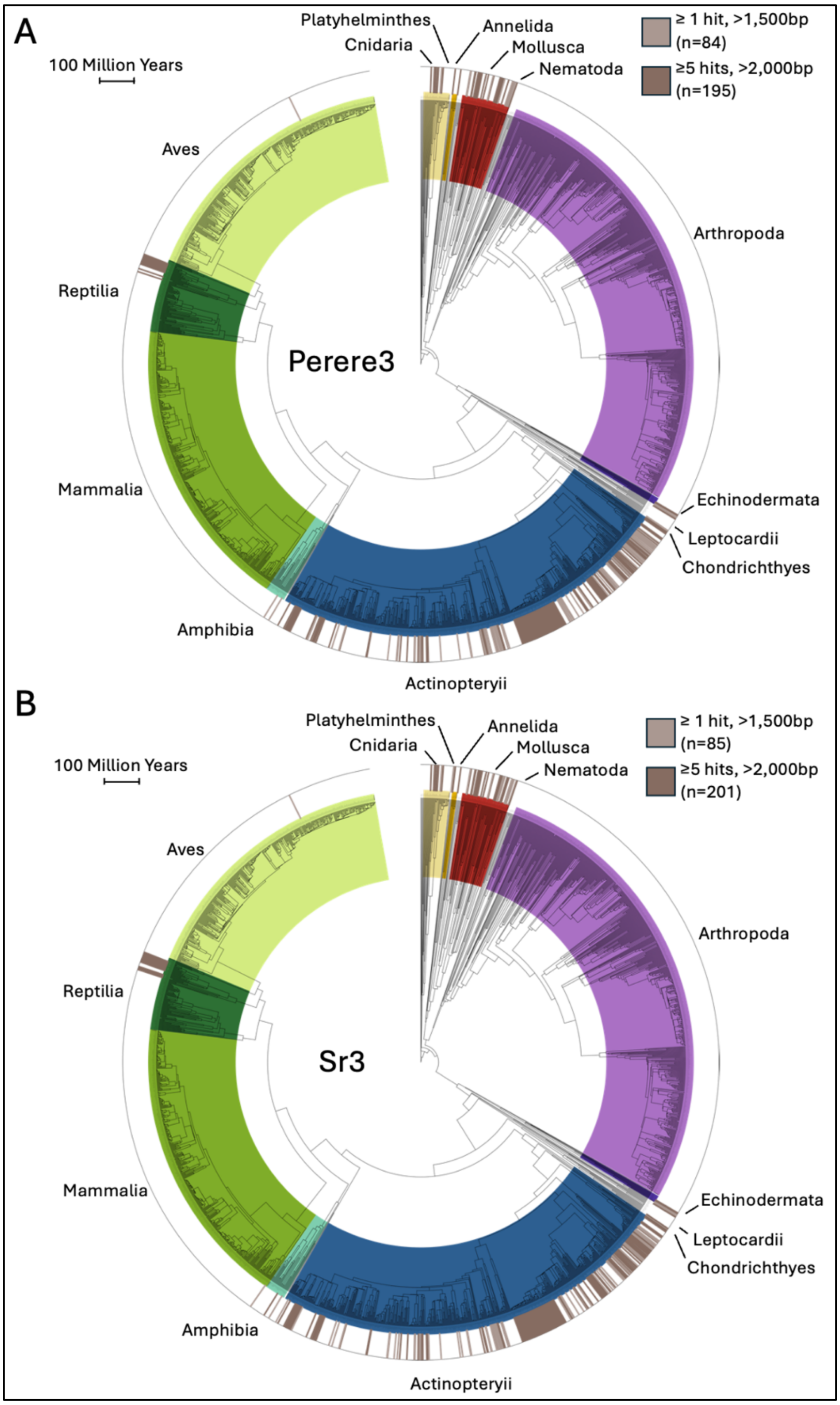


**Supplementary Figure 1 - Separate Perere-3 (A) and Sr3 (B) blast results across high-quality metazoan genomes.** Presence of Perere-3 / Sr3-like sequences across metazoan genomes available at Timetree.org. Relevant hits were classified as either weak (light brown: at least one hit > 1,500bp) or strong (dark brown: at least one hit > 2,000bp and at least five hits > 1,000bp).
